# Inhibiting Bet1-mediated transport of MMP14 to plasma membrane impaired GBM cell invasion

**DOI:** 10.1101/2023.02.16.525994

**Authors:** Yani Luo, Jiana Li, Yi Xiong

## Abstract

**Purpose:** Glioblastoma (GBM) is the most aggressive and common form of brain cancer in adults. GBM is characterised by poor survival as the lack of effective therapies. This research aims to detect the roles of SNAREs in GBM and improve our knowledge of targeting therapy for GBM.

**Materials and methods:** the expression of SNAREs and their correlation with overall survival (OS) in GBM are investigated using the GEPIA. The level of BET1 in GBM cell lines was tested by RT-qPCR, and its biological functions in GBM cells were tested by Transwell assay and CCK8 kit. The effect of BET1 on the location of MMP14 is identified by Immunofluorescence.

**Results:** The expression profile of SNARE family members in GBM tissue is changed dramatically. Among them, the mRNA levels of BET1 and VAMP3 are up-regulated, and their expression negatively correlates with OS. BET1 is also increased in GBM Cell Lines, and it is required for efficient GBM cell migration and invasion partly because it mediates the transport of MMP14 to the plasma membrane.

**Conclusion:** GBM has highly diffusive and infiltrative ability in nature, making complete surgical resection almost impossible. Our data shows that BET1 is highly expressed in GBM tissue, negatively correlated with OS, and essential for GBM cell migration and invasion. These results indicate that SNARE BET1 may present a potential target for GBM treatment.

## Introduction

Glioblastoma (GBM) originating from astrocytes, neural stem cells or progenitors is one of the most aggressive solid tumours.^1^ The clinical treatment involves maximal surgical resection added with adjuvant temozolomide therapy and radiotherapy.^2^ However, despite many efforts that have been made, the 5-year survival rate of GBM patients is only 5%.^3^ Thus, knowledge of the underlying mechanisms of GBM is urgently needed to develop effective therapies.

SNAREs have more than 39 family members that mediate membrane fusion of target membranes with transport vesicles.^4^ The cargos of vesicles include membrane proteins, lipids, signalling molecules, hydrolytic enzymes and biosynthetic, and the trafficking machinery itself. The proper function of membrane trafficking is required for cell-cell communication, division, cellular growth, division and movement.^5^ These biological process is essential for GBM growth and metastasis.

Several SNAREs have been reported to play significant roles in the progress of GBM. Blockade of STX1 inhibited GBM cell proliferation, invasion, and tumorigenicity after grafting GBM cells into the brain of mice. ^6^ The STX17-SNAP29-VAMP8 SNARE complex is involved in the radiosensitivity of GBM cells by mediating the fusion between autophagosomes and lysosomes.^7^ Mig-6 mitigates the malignant potential of GBM cells by dampening STX8-mediated transport of EGFR and driving EGFR into late endosome and lysosome to degradation.^8^ While SNAP25 inhibits GBM progression by regulating synapse plasticity by GLS-mediated glutaminolysis.^9^

However, the roles of other SNARE family members are poorly illustrated in GBM. In this study, we first investigated the expression of SNAREs in GBM compared to normal tissue and then the correlations between the level of SNAREs with patients’ overall survival. Finally, we tested the biological function of ER-Golgi SNARE BET1 in GBM cells and the related mechanism.

## Material and methods

### Gene Expression Analysis and Overall Survival Analysis

The differentially expressed SNAREs in GBM tissues compared to normal tissues were identified using the GEPIA (Gene Expression Profiling Interactive Analysis, http://gep.cancer-pku.cn/index.html). ^10^ The mRNA-seq data for TCGA GBM were matched with normal TCGA and GTEx data. Collected data were shown as transcripts per million (TPM) values after transformation, and the differential expression analysed by boxplot was performed using log2 (TPM+1) values. 163 biopsies from patients with GBM and 207 with normal tissue were used in this analysis.

Overall Survival (OS) analyses were conducted using the GEPIA. Survival curves were estimated using Kaplan–Meier estimator. Survival curves were compared by the log-rank test. 161 GBM patients were analysed from the TCGA dataset.

### Cell Culture and Cell proliferation test

Glioma cell lines U251, U-118 MG (gifts from The Hong Kong University of Science and Technology), A172, U-87 MG (Procell, Wuhan, China), and human astrocyte cell line HA1800 (Jennio Biotech, Guangzhou, China) are cultured according to TACC suggestion. The cells were cultured with 5% CO2 at 37°C.

CCK-8 kit (Beyotime, Shanghai, China) was used to measure the proliferation of U-87MG and A172 cells. 1500 cells per well were cultured in 96-well plates in the complete MEM medium. CCK-8 reagent (10 μL) was added to 90 μL MEM to generate a working solution and incubated for 2 h after transfection. We performed CCK-8 assay after transfection 0 h, 24 h, 48 h, 72 h, and 96 h.

### siRNAs, Plasmid constructs, and transfection

Total RNA extracted from A172 cells was used to reverse-transcription reaction and synthesise complementary DNA, and the ORF encoding MMP14 was then PCR-amplified using its primers. The amplified ORF DNAs were cloned into pCMV-FLAG. The siRNA targeting BET1 is siRNA#, 5’-AAGCAAAGTAACTGCTATAAA-3’; siRNAs, Plasmids and their negative controls were transfected with RNAiMAX and Lipofectamine 2000 (Thermo Fisher Scientific), respectively, as the manufacturer’s protocol. The knockdown efficiency of siBET1 is in supplements.

### RT-qPCR

Total RNA was extracted from the cell lines using the TRIzol RNA Isolation Reagents (sigma, St. Louis, MO, USA). The first-strand cDNA was converted using a reverse transcription system (Promega, Wisconsin, USA) according to the manufacturer’s instructions. RT-qPCR analysis was performed using iTaq TM Universal SYBR Green Supermix (Bio-Rad, California, USA). GAPDH was used for normalisation. Primaries are shown in Table 1.

**Table 1.**
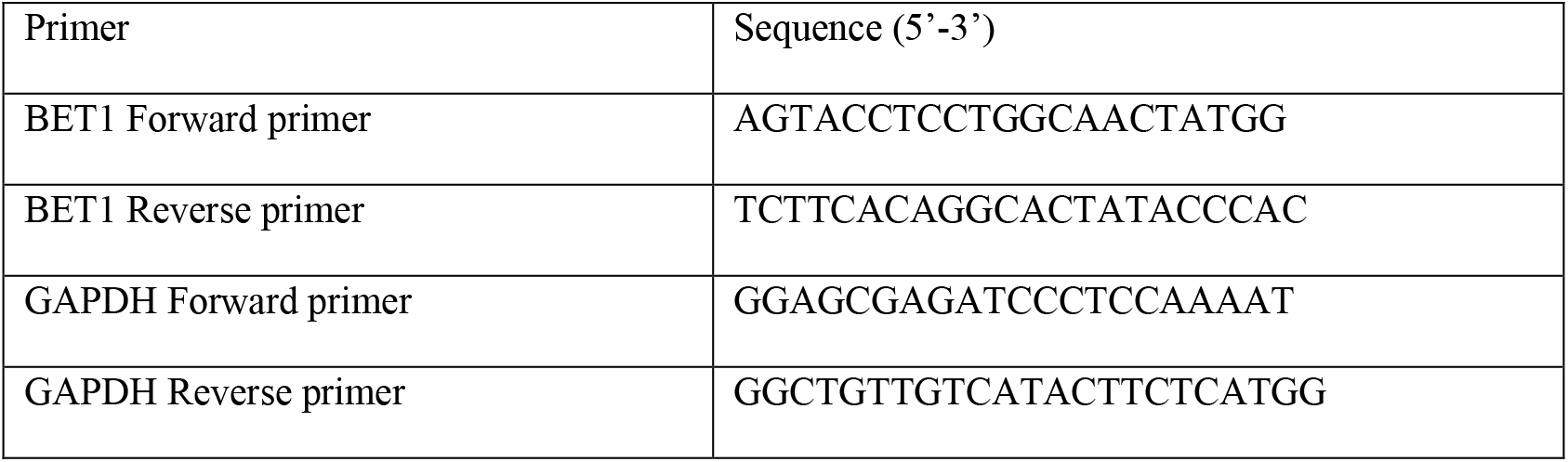

### Transwell invasion and migration assay

Transwell chamber (Corning, New York, USA) with 8-μm pores was used to test the ability of cell invasion and migration. After cells transfected with siBET1 or its negative control (NC) for 48h, for the invasion test, cells were seeded into chambers coated with Matrigel (BD) with serum-free MEM medium and bottom chambers with complete medium with 10% FBS as a chemoattractant, and cultured for 24 h. For the migration test, cells were seeded in chambers without Matrigel and cultured for 12 h under the same condition as the invasion test. The invaded and migrated cells were stained with crystal violet and pictures were captured by microscopy.

### Immunofluorescence

Immunostaining and confocal microscopy were performed as described previously^11^. Briefly, cells were cultured on glass coverslips coated with gelatin, fixed by 4% paraformaldehyde, permeabilised by 0.1% Triton X-100, and then stained with the antibodies of MMP14 and DAPI. To detect cell plasma membrane MMP14 (PM-MMP14), cells were incubated with antibodies that reacted with the extracellular domain of MT1-MMP (R&D Systems, Minnesota, USA) in the complete medium for 30 min on ice before fixation and permeabilisation. CarlZeiss LSM710 laser-scanning confocal microscope was used to capture images. The mean fluorescence intensity (MFI) of PM-MMP14 was analysed with the Image J Software (National Institutes of Health, Bethesda, USA).

### Statistics

All statistical analyses were performed with GraphPad Prism 6.01 software. Significant differences (P<0.05) between the two groups were identified by Student’s t-tests.

## Results

### The expression profiles of SNARE family members in GBM

The expression of SNARE family members was analysed in biopsies from patients with GBM and normal biopsies using the GEPIA.^10^ As figure1 shows, the expressions of BET1 STX8 STX10 SEC22A YKT6 STX11 SNAP23 VAMP7 VAMP3 BNIP1 VAMP8 VAMP5 and SNAP29 are increased in GBM, while SNAP25, STX1B, VAMP1 and VAMP2 are decreased in GBM. The expression level of the genes of interest was obtained in 207 samples from normal tissue and 163 from patients with GBM.

**Figure 1.**
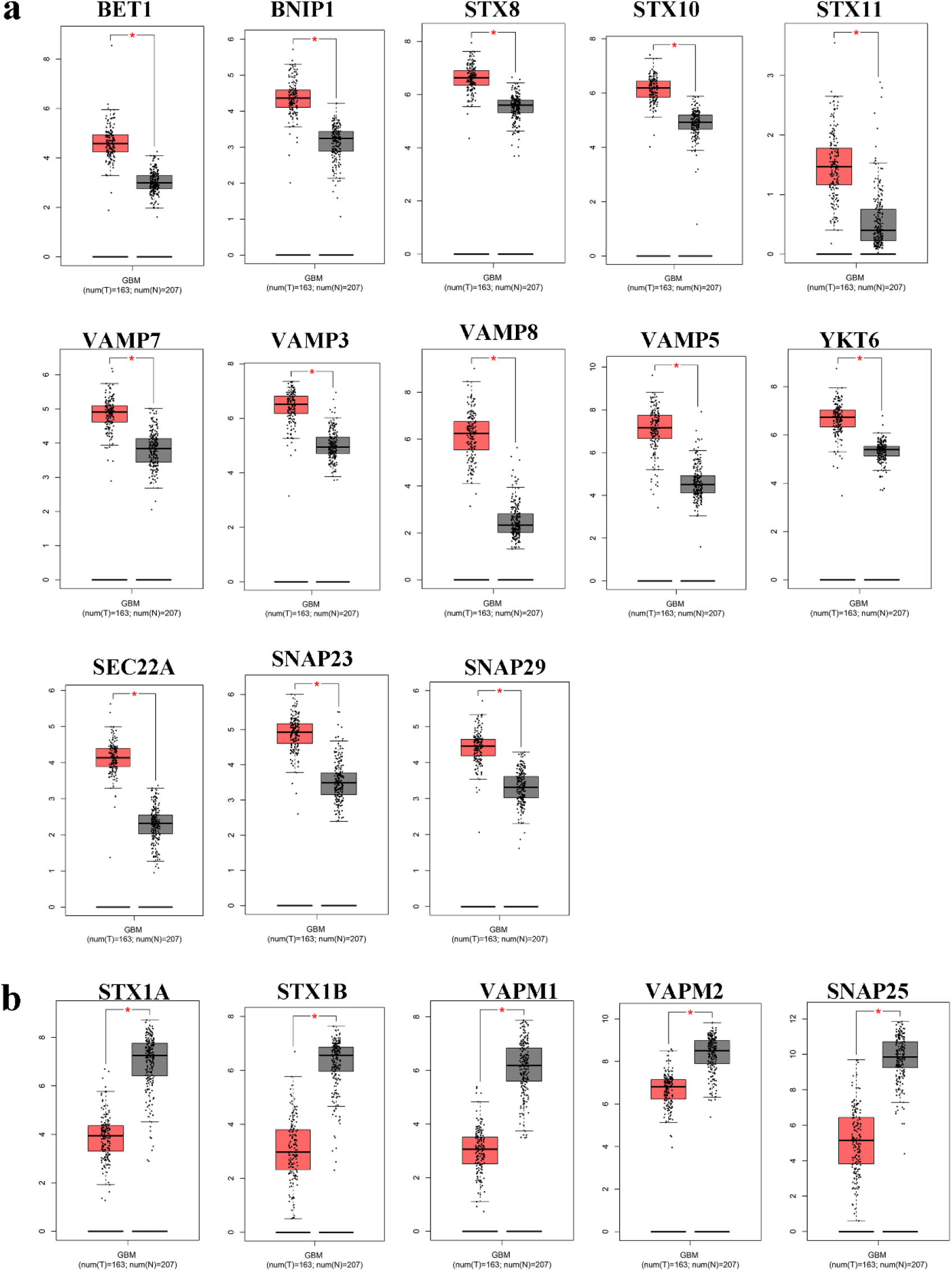
Differentially expressed SNAREs in GBM tissue. (a) represents up-regulated SNAREs in biopsies of patients with GBM (T) (red box) compared with normal tissue (N) (grey box), including BET1 STX8 STX10 SEC22A YKT6 STX11 SNAP23 VAMP7 VAMP3 BNIP1 VAMP8 VAMP5 and SNAP29. (b) shows down-regulated SNAREs in biopsies of patients with GBM (T) (red box) compared with normal tissue (N) (grey box), including SNAP25, STX1B, VAMP1, and VAMP2. Data were obtained from the TCGA and GTEx dataset. * p < 0.01. Expression was log2 transformed (TPM+1).

### High Expression of BET1 and VAMP3 Correlates with Lower Overall Survival in patients with GBM

To investigate whether differentially expressed SNAREs are involved in survival in patients with GBM, OS analyses were performed using Kaplan–Meier estimator. As figure 2 shows, higher expression of BET1 and VAMP3 genes were significantly associated with shorter OS in patients diagnosed with GBM with hazard ratios of 1.8 and 1.6, respectively.

**Figure 2.**
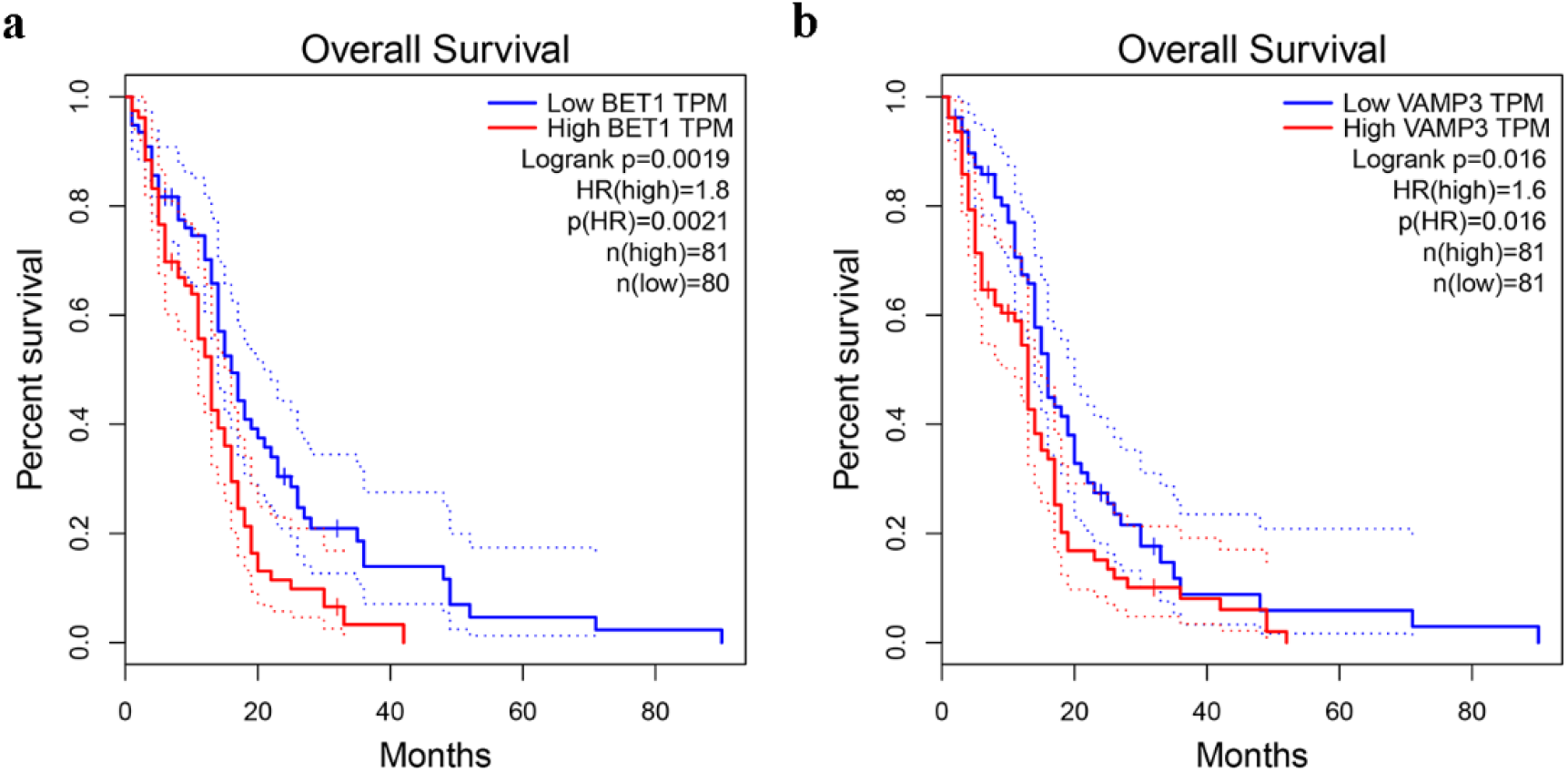
Kaplan–Meier curves of overall survival in GBM patients by level of BET1 (a) and VAMP3 (b). The results indicate the probability of survival for 80 months. The lines in red show high expression, and the blue colour shows the low expression of genes. p < 0.05 were regarded as statistically significant.

### BET1 Is Increased in GBM Cell Lines and Required for efficient GBM cells’ migration and invasion

To determine whether, as in patient tissues, the expression of BET1 is up-regulated in GBM cell lines, its expression was tested in the GBM cell lines U-87MG, U-118MG, A172, and U251 by RT-qPCR. The results show that the mRNA expression level of BET1 increased in four GBM cell lines compared to the human astrocyte cell line HA1800 (Figure 3a). One of the reasons for the high mortality is that GBM has highly diffusive and infiltrative ability in nature, making complete surgical resection almost impossible.^12^ We have conducted the Transwell invasion and migration assay to test the role of BET1 in GBM cells. As shown in figure 3b-c, knockdown BET1 significantly inhibited the migration and invasion of U118, A172, and U251 cells. However, down-regulating the level of BET1 did not significantly influence U-87MG cells’ migration and invasion (data not shown).

**Figure 3.**
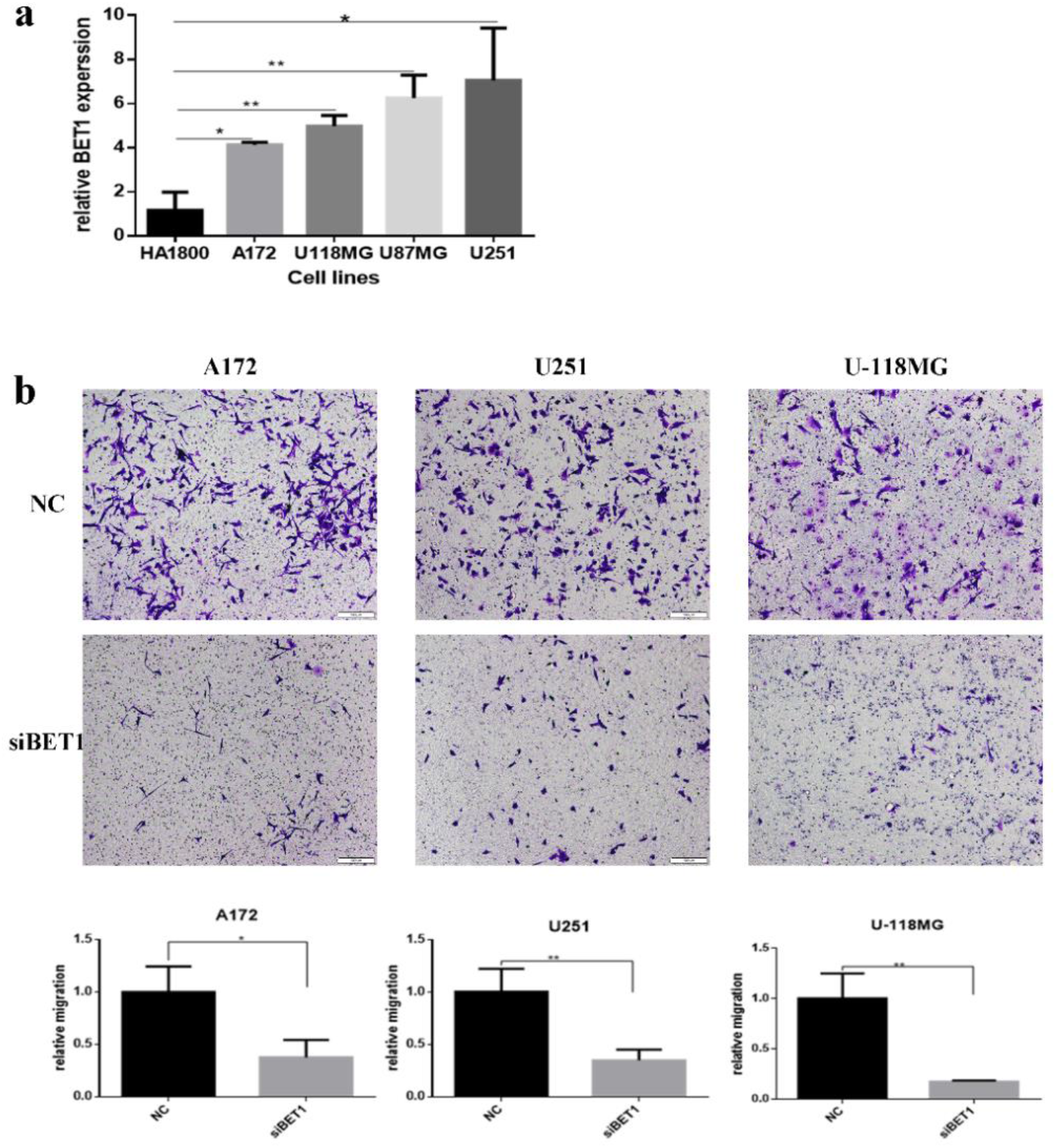
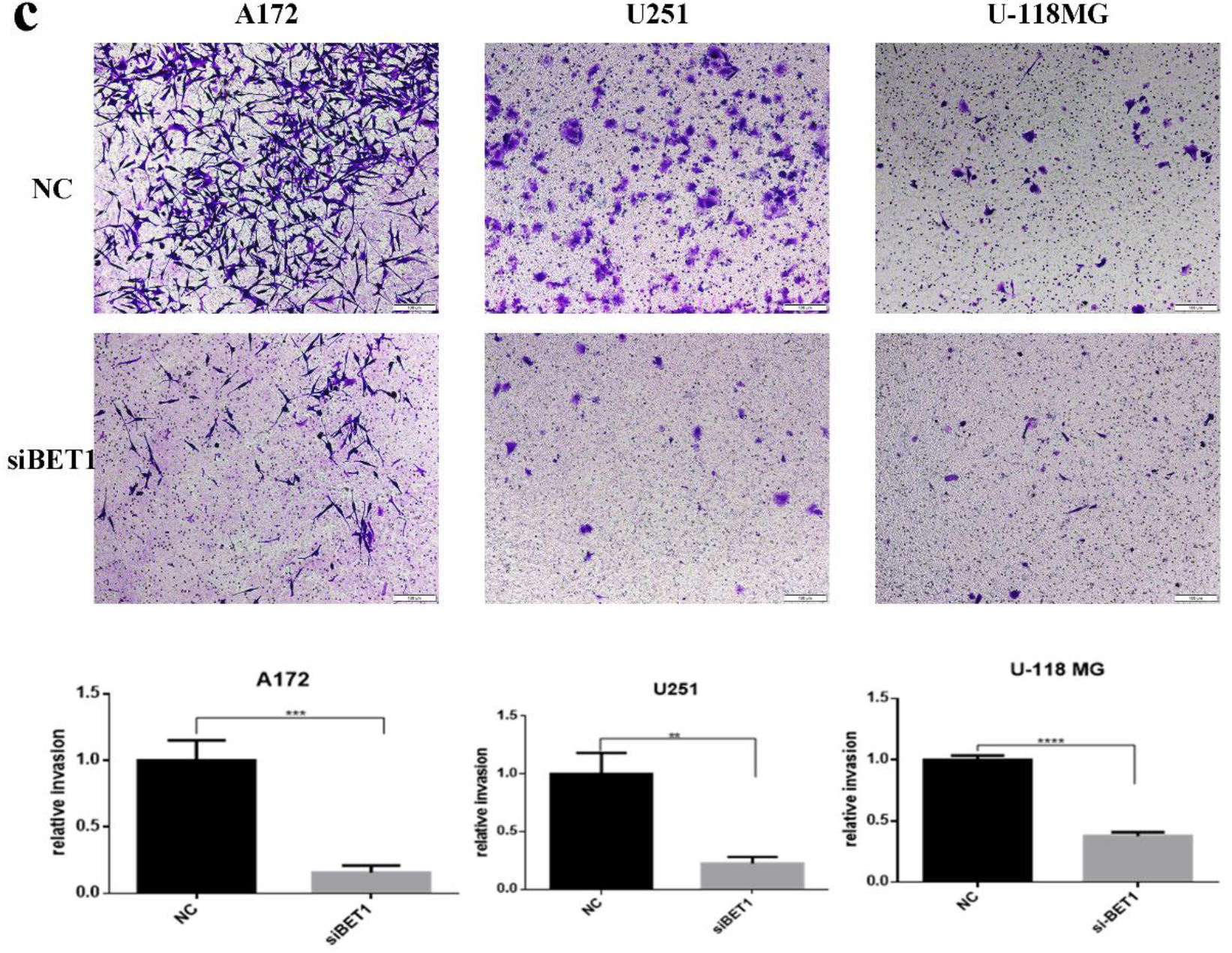
BET1 is required for effective GBM cell migration and invasion. (a) the relative mRNA expression of BET1 in glioma cell lines A172, U-118MG, U-87MG and U251 compared to human astrocyte cell line HA1800. (b) Representative micrographs and statistical data of Transwell migration assay. Transwell migration assay performed in A172, U251, and U-118MG transfected with siBET1 or NC. (c) Representative micrographs and statistical data of Transwell invasion assay. Transwell invasion assay performed in A172, U251, and U-118MG transfected with siBET1 or NC. (Scale bar: 50 μm; *p<0.05, **p<0.01, ***p<0.0001, n=3)

### MMP2, MMP9, and MMP14 Expression Is Up-regulated in Samples from Patients with GBM

Remodelling the extracellular matrix, which depends on the secreted MMPs (MMP2, MMP9) and membrane-bound membrane-type–MMPs (MMP14), is essential for cell invasion and metastasis.^13^ Our database mining results show that MMP2, MMP9, and MMP14 are all highly expressed in GBM compared to normal tissues (Figure 4a), consistent with previous reports.^14–16^ In this study, we focused on MMP14 because it is an activator of MMP2 ^17^ and essential for GBM cell migration and invasion.^18^ Also, its expression negatively correlates with GBM patients’ overall survival (Figure 4b).

**Figure 4.**
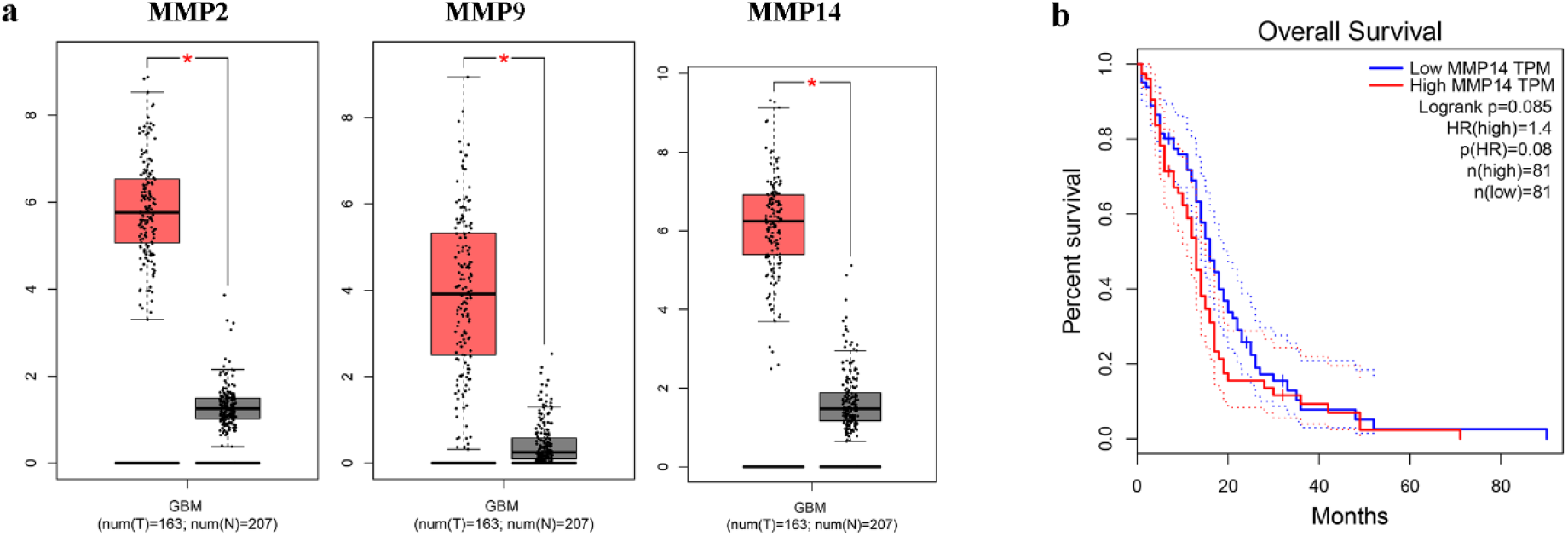
MMP2, MMP9, and MMP14 are increased in GBM, and MMP14 significantly correlates with patients’ OS. (a) Expressions of MMP2, MMP9, and MMP14 in tissues of GBM patients (T) (red box) and normal tissue (N) (grey box) using the GEPIA. * p < 0.01. (b) Kaplan–Meier curve of overall survival in GBM patients by level of MMP14. The lines in red show high expression, and the blue colour shows the low expression of MMP14. p < 0.05 were regarded as statistically significant.

### Knocking down the level of BET1 impaired the transportation of MMP14 to the plasma membrane

MMP14 requires transport to the plasma membrane via trafficking vesicles for action. ^19,20^ We use an antibody which reacts with the extracellular domain of MMP14 before permeabilisation to detect the level of MMP14 located at the plasma membrane in A172 cells. BET1 depletion decreased the MMP14 level on the plasma membrane (Figure 5a-b). This result is consistent with the total MMP14 distribution. BET1 depletion causes the total MMP14 to centralised around the nuclear, while in the control cells, the MMP14 are distributed evenly (Figure 5c). These results indicate that Bet1 participates in the delivery of MMP14 to the plasma membrane.

**Figure 5.**
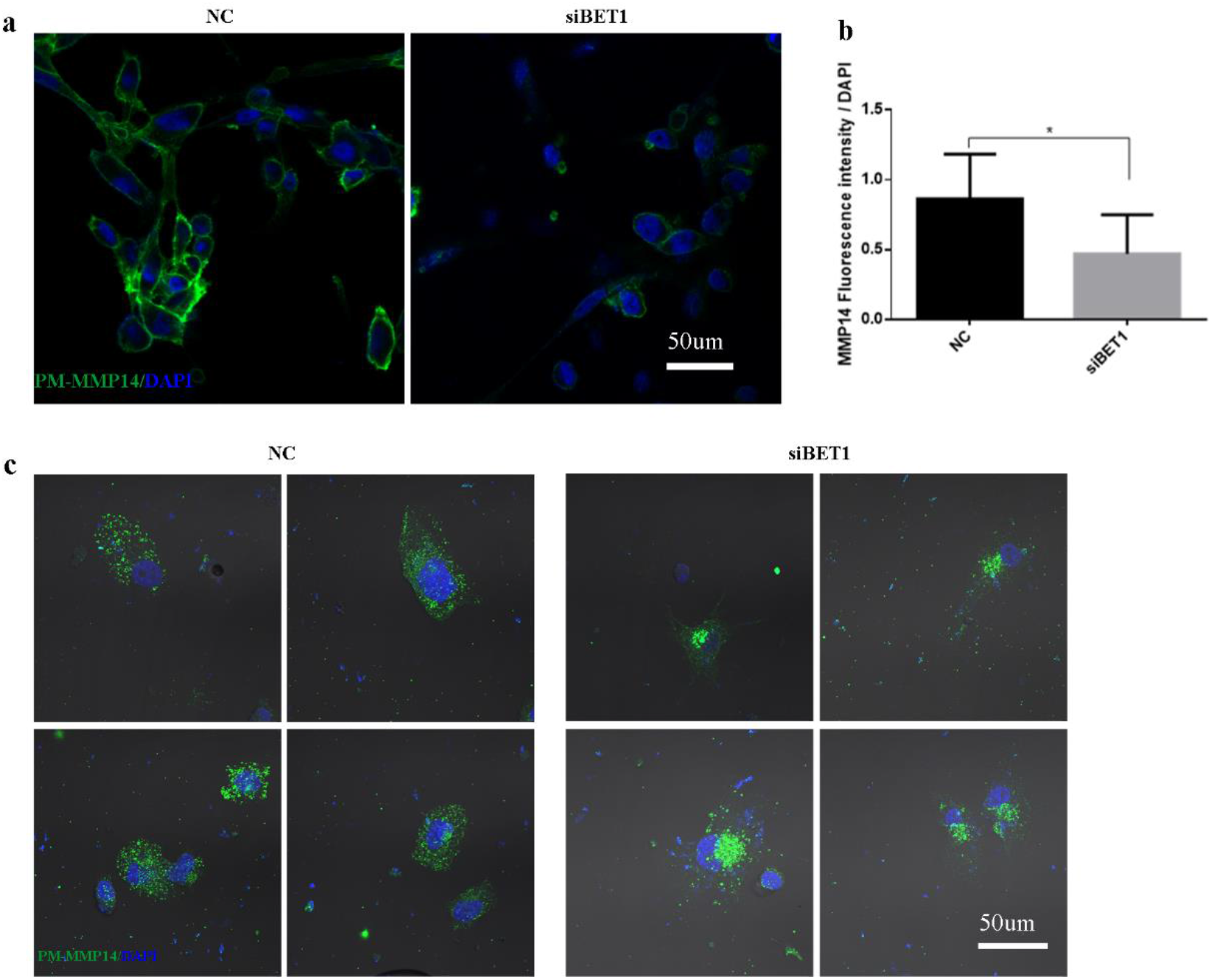
BET1 mediated MMP14 transport to the plasma membrane. (a-b) immunofluorescence of representative image of the intensity of PM-MMP14 (a) (left for negative control A172 cells, right for siBET1-treated A172 cells), nuclei stained with DAPI, PM-MMP14 stained with green and quantitative data (b). (c) representative images of the distribution of total-MMP14 (left for negative control A172 cells, right for siBET1-treated A172 cells), nuclei stained with DAPI, and total-MMP14 in green. (Scale bar: 50 μm; *p<0.05, **p<0.01, ***p<0.0001, n=3)

### Knocking down the level of BET1 inhibited the proliferation of GBM cells

Bet1 is a protein involved in the transport from the Endoplasmic reticulum to the Golgi body, a vital process for cell functions. Consistent with this, BET1 knockdown significantly inhibited the proliferation of GBM cell lines U-87MG and A172 tested by CCK8 kit (Figure 6).

**Figure 6.**
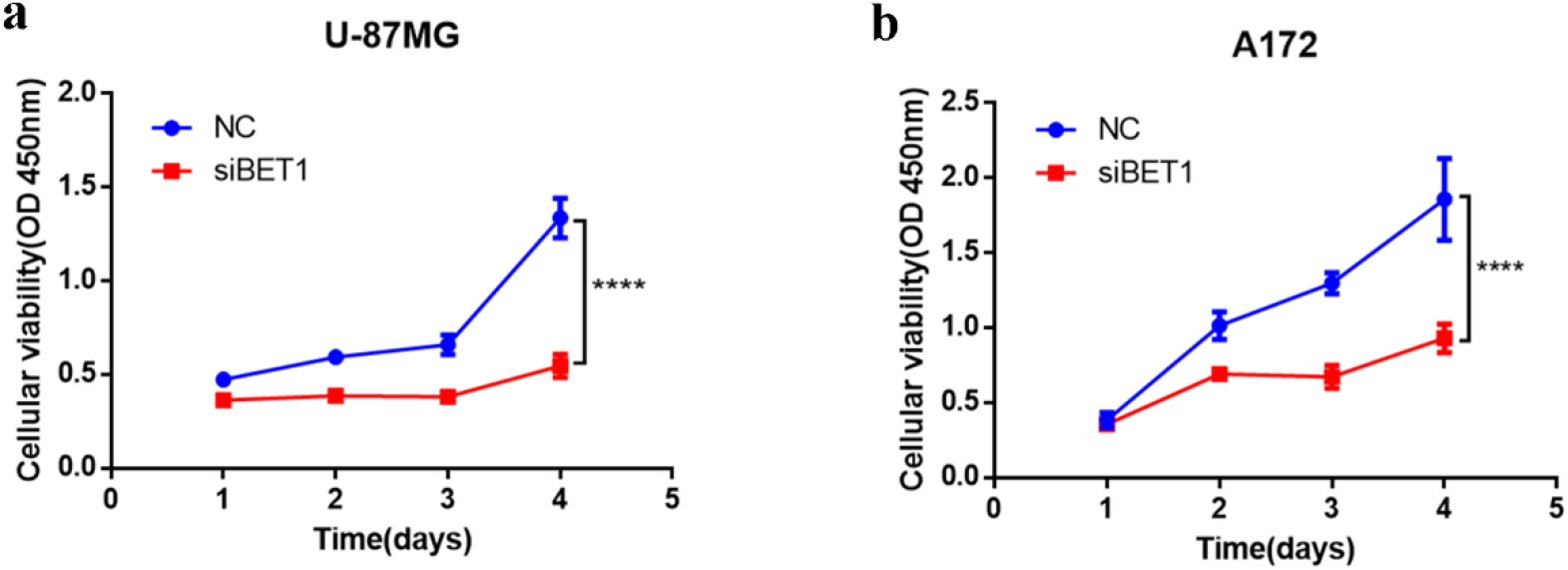
Lower expression of BET1 inhibited glioma cell line proliferation. CCK-8 kit was used to detect the proliferation of U-87MG (a) and A172 (b) at 1, 2, 3, and 4 days after transfection of siBET1 or NC.

## Discussion

Many tumour-promoting proteins are active spatiotemporally. the underlying mechanisms involve their trafficking to and from specific subcellular sites, which depends on SNARE proteins mediated membrane fusion.^5^ We found that the expressions of SNARE family members change dramatically in GBM patients compared with normal tissues by mining the GEPIA dataset. STX8 involved in EGFR trafficking and gliomagenesis^8^ and SNAP29-VAMP8 mediating autophagy in radioresistant GBM cells ^7^ are increased in GBM tissues compared to normal tissues, while a reported GBM suppressor SNAP25 that inhibits GBM cell proliferation, migration, invasion and fostered metabolism reprogramming is decreased. ^9^ However, our results show that STX1a and STX1b are significantly decreased in the GBM sample. It seems to conflict with a previous report that STX1 is required for GBM cell line U373 proliferation and invasion.^6^ The reasons for this need more exploration. The roles of other differentially expressed SNARE genes in GBM, especially VAMP3 and BET1 that are significantly correlated with GBM patients’ OS, are short of illustration and need to be further investigated.

In this study, we found that BET1 is essential for GBM cells invasion and migration, partly because it mediates the transport of MMP14 to the plasma membrane, where MMP14 promotes cell metastasis by focal pericellular degradation of the ECM.^18^

Pro-MMP14 is synthesized in ER and obtains catalytical activity in the trans-Golgi network (TGN) through furin-mediated proteolytic cleavage before its arrives at the plasma membrane. ^21,22^ This process relies on COPII vesicle transport mediated by the STX5-GS28-Ykt6-Bet1 complex.^23^ However, the mechanism of BET1-mediated MMP14 transport may be more complicated. Endocytic recycling is a way to optimize the proteolytic activity of MMP14 on ECM degradation.^24^ Late endosome is a major compartment for intracellular MMP14 storage.^25^ A previously reported study shows that BET1 is not only localized and functions in the ER-Golgi secretory pathway but also mediates the transport from MMP14-positive late endosomes to the plasma membrane in invasive breast cancer cell line MDA-MB-231. But this process does not exist universally in all cells, such as not in Hale cells. ^26^ It requires further investigation to test whether BET1 -mediated transport of MMP14 transport in GBM cells is involved in the transport of MMP14-positive late endosomes. COPII vesicle-mediated ER-Golgi transport is vital for cell survival and proliferation. correspondingly, our results show that the low level of BET1 inhibited GBM cell proliferation.

## Conclusion

GBM has highly diffusive and infiltrative ability in nature, making complete surgical resection almost impossible.^12^ Our data show that BET1 is highly expressed in GBM tissue, negatively correlated with patients’ OS, and essential for GBM cell migration and invasion. These results indicate that SNARE BET1 may present a potential target for GBM treatment.

