## Supplementary figures and images for "Inhibiting Bet1-mediated transport of MMP14 to plasma membrane impaired GBM cell invasion"

### supplements

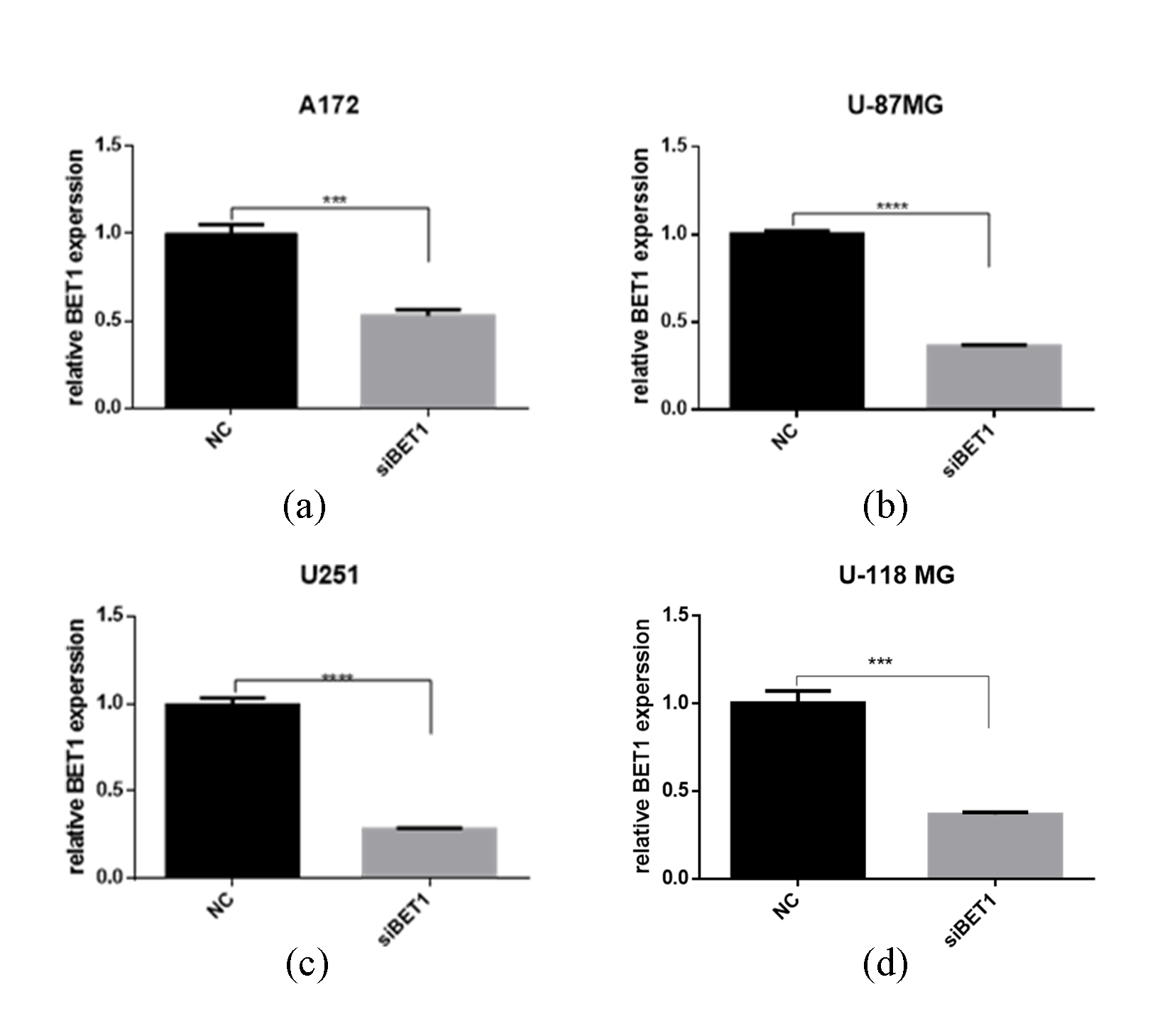
